# Targeting USP2 induces degradation of PML-RARα with or without drug-resistant mutations in acute promyelocytic leukemia

**DOI:** 10.1101/2025.05.11.653221

**Authors:** Jie Zhang, Wenxuan Wu, Yun Wang, Youping Zhang, Yingying Wang, Wenhui Bai, Zhenge Zhang, Chujiao Zhu, Yunzhao Wu, Ziwei Zhang, Li Yang, Hu Lei, Hanzhang Xu, Li Zhou, Yingli Wu

## Abstract

Despite the high efficacy of all-trans retinoic acid (ATRA) and arsenic trioxide (ATO) in treating acute promyelocytic leukemia (APL), approximately 10-20% of patients develop drug resistance due to mutations in PML-RARα and other factors. Here, it is demonstrated that inhibiting USP2 with ML364 or silencing USP2 reduces PML-RARα protein levels in both ATRA-sensitive and -resistant APL cells, an effect reversed by proteasome inhibition. Conversely, USP2 overexpression enhances PML-RARα stability. Mechanistically, USP2 interacts with and deubiquitinates PML-RARα, including drug-resistant mutants. Consistent with PML-RARα degradation, ML364 treatment significantly induces apoptosis in APL cell lines and primary leukemia cells. In conclusion, this study identifies USP2 as a novel deubiquitinating enzyme for PML-RARα and highlights USP2 inhibition as a potential therapeutic strategy for APL with PML-RARα mutations.

## Introduction

PML-RARα is generated by t(15;17)(q24;q21) chromosomal translocation[1]. The resulting oncoprotein blocks cell differentiation and suppresses apoptosis, ultimately leading to the development of acute promyelocytic leukemia (APL) [2, 3]. The introduction of all-trans retinoic acid (ATRA) and arsenic trioxide (ATO) has significantly improved the overall survival of APL patients [1, 4, 5]. However, approximately 10-20% of APL patients develop resistant to the treatment with ATRA and/or ATO[6]. Among the known resistance mechanisms, the most prevalent involve mutations in the PML-RARα fusion gene, particularly in the ligand-binding domain (LBD) of RARα and the PML-B2 domain, the primary target site of ATO [7–9]. Mutations in the PML-B2 domain (e.g., A216V and L218P) may impair ATO therapeutic efficacy by disrupting proper ATO binding, interfering with post-translational modifications (such as SUMOylation and multimerization), and causing aberrant subcellular localization [10–12]. These defects collectively lead to insufficient degradation and pathological retention of oncoproteins, ultimately sustaining their stability and driving therapeutic resistance by evading ATO-mediated clearance mechanisms. When mutations occur in LBD of RARα, conformational changes can weaken its binding affinity to ATRA. Due to this reduced binding capacity, the LBD may fail to properly dissociate nuclear receptor corepressor complexes and recruit activator complexes, leading to aberrant transcriptional regulation and subsequent ATRA resistance [13–15]. One promising strategy to overcome this mutation-induced drug resistance is to induce the protein degradation by alternative pathways. For example, targeting HDAC3 has been shown to induce the degradation of PML-RARα in ATRA- and ATO-resistant APLs[16].

Deubiquitinases (DUBs) regulate protein stability and function by removing ubiquitin chains from substrate proteins, modifying ubiquitin linkages, and processing ubiquitin precursors [17]. The stability of PML-RARα can also be influenced by DUB activity, suggesting that inhibiting specific DUBs may induce PML-RARα degradation and overcome ATRA- and ATO-resistance in patients with PML-RARα mutations. For example, inhibition of YOD1, a member of the varian tumor protease (OTU) family, has been shown to promote PML-RARα degradation and effectively eliminate APL cells, including drug-resistant subtypes [18]. This work suggests that DUB inhibition as a potential therapeutic strategy for degrading PML-RARα.

In the present study, we identified USP2 as a novel DUB for PML-RARα. Knockdown USP2 or use of the USP2 inhibitor ML364 could induce degradation of PML-RARα and apoptosis in APL cells. Importantly, inhibition of USP2 could also degrade drug-resistant PML-RARα mutants. Our study demonstrated that USP2 is a novel target for the treatment of APL, particularly in cases of drug-resistant APL driven by PML-RARα mutations.

## Materials and methods

### Cells and culture

Human embryonic kidney (HEK293T) cells were purchased from the American Type Culture Collection (ATCC, Manassas, VA, USA). NB4 and MR2 cells were provided by Prof. Michel Lanotte (Hôpital Saint-Louis, Paris, France). NB4 and MR2 cells were cultured in RPMI-1640 medium (BasalMedia, Shanghai, China). HEK293T cells were cultured in Dulbecco’s modified Eagle’s medium (DMEM; BasalMedia, Shanghai, China). All of the media were supplemented with 10% fetal bovine serum (FBS; Vazyme, Nanjing, China) and 1% penicillin/streptomycin (BasalMedia, Shanghai, China). All cell lines were maintained at 37 °C in a humidified atmosphere containing 5% CO_2_. Primary APL blasts extracted from the bone marrow of APL patients (Ruijin Hospital, Shanghai Jiaotong University School of Medicine, Shanghai, China) were isolated using Lymphoprep™ (STEMCELL, Vancouver, Canada). Written informed consents from patients and approval from the Institutional Review Board of Ruijin Hospital, Shanghai Jiao Tong University School of Medicine were obtained before the use of these clinical materials for research purposes.

### Plasmids, reagents and antibodies

The PML-RARα plasmid was a kind gift from Prof. Kan-Kan Wang (Shanghai Institute of Hematology, Rui Jin Hospital, Shanghai Jiao Tong University School of Medicine). PML-RARα drug-resistant mutants (A216V, L218P, R276Q and ΔF286) were kind gifts from Prof. Jiong Hu (Rui Jin Hospital, Shanghai Jiao Tong University School of Medicine). PLZF-RARα plasmid was preserved in our laboratory. USP2^WT^ (USP2 wild-type), USP2^C276S^ (USP2 catalytic mutant), USP2 (1-258 amino acids) and USP2 (259-605 amino acids) plasmids were constructed in our lab. The HA-USP8 plasmid was purchased from YouBio (Hunan, China). The USP2 shRNAs were purchased from Tsingke Biotechnology Co., Ltd. (Shanghai, China). USP2 shRNA-1 and USP2 shRNA-2 sequences: USP2 shRNA #1, 5′-GCTACACAGATGCCCACTATG-3′; USP2 shRNA #2, 5′-GGAGTTCCTTCGCTTTCTTCT-3′.

ML364, b-AP15, Degrasyn and USP25/28 inhibitor AZ1 were purchased from MedChemExpress. P22077 was purchased from EMD Millipore. ATRA, ATO, MG132, CQ, and cycloheximide (CHX) were purchased from Sigma–Aldrich.

Antibodies against RARα (#62294) were purchased from Cell Signaling Technology. Antibodies against PML (sc-71910) were purchased from Santa Cruz. Antibodies against USP2 (10392-1-AP) and β-actin (66009-1-Ig) were purchased from Proteintech. Antibodies against Myc (AE070), HA (AE105) and Flag (AE063) were purchased from ABclonal. Antibodies against His (ab18184) were purchased from Abcam.

### Lentivirus production and transduction

HEK293T cells were co-transfected with transfer plasmid and packaging plasmids (psPAX2 and pMD2.g). After transfected for 6 hours, old medium was replaced with fresh pre-warmed complete medium. After another 48 hours, viral particles were collected and resuspended in RPMI-1640 medium and then frozen at −80L°C. NB4 and MR2 cells were infected with the virus in the presence of polybrene (8 μg/mL) to enhance the infection efficiency. After 3 days of infection, cells were screened by puromycin (1 μg/mL) for at least 5 days.

### Immunoprecipitation and western blotting

For immunoprecipitation assays, cells were lysed in lysis buffer (20 mM Tris-HCl, pH 7.5; 150 mM NaCl; 0.1 mM EDTA; 0.2% Triton X-100) supplemented with protease inhibitor cocktail (MedChemExpress, Monmouth Junction, NJ, USA) on ice for 30 minutes. After 13,000 g centrifugation for 15Lminutes at 4 °C, the supernatants were incubated with primary antibodies at 4 °C overnight. Protein A/G agarose beads (Beyotime, Shanghai, China) were added for 5-6 hours the next day. Next, the beads were washed with the washing buffer (20 mM Tris-HCl, pH 7.5; 150 mM NaCl; 0.1 mM EDTA; 1% Triton X-100) three times. Finally, the bound proteins were dissolved in 5× SDS-PAGE sample loading buffer and analyzed by western blotting. For ubiquitylation analysis, MG132 (15 μM) was added 4-6 hours before cell collection.

After the corresponding treatment, cells were collected and lysed with 2× SDS sample loading buffer. Protein extracts were electrophoresed using 6%-10% SDS-PAGE gels (Meilunbio®, Liaoning, China) and transferred to nitrocellulose membranes (Bio-Rad, Hercules, CA, USA). Membranes were blocked with 5% non-fat milk at room temperature for 1 hour and incubated with primary antibodies overnight at 4 °C. The membranes were then incubated with the horseradish peroxidase (HRP)-conjugated secondary IgG antibody at room temperature for 1 hour. The membranes were finally imaged by chemiluminescence (ECL, Amersham, Little Chalfont, UK).

### Cell proliferation and cell viability assay

To assess cell viability, cells were collected and counted using the Countstar automated cell counter. Before counting, trypan blue was used to distinguish between viable and dead cells.

Cell proliferation assay was performed using the Cell Counting Kit-8 (Meilunbio®, Liaoning, China). Cells were cultured in 96-well or 384-well plates and then treated with appropriate concentrations of drugs. After incubation for 2-3 days, CCK-8 solution was added to each well for 2-4 hours and the optical density (OD) at 450 nm was detected using a microplate reader.

### Cell apoptosis and differentiation analysis

Apoptosis of cells was detected using Annexin V and 7-Aminoactinomycin D (7-AAD) or propidium iodide (PI) staining. After treatment, the cells were collected and washed with PBS buffer and then stained according to the manufacturer’s protocol. Early apoptotic cells (Annexin V-positive, 7-AAD or PI-negative) and late apoptotic cells (Annexin V-positive, 7-AAD or PI-positive) were then determined by flow cytometer (BD Biosciences, San Diego, CA, USA). The results were analyzed using CytExpert 2.4 software. The kits include 488-Annexin V and PI Apoptosis Kit (Green), 647A-Annexin V and PI Apoptosis Kit (Red) (SB-Y6002, SB-Y6026, ShareBio, Shanghai, China) and Annexin V-APC/7-AAD Apoptosis Kit (AP105, Liankebio, Hangzhou, China).

Cell differentiation was evaluated by the expression of cell surface maker of differentiation. After treated with ATRA and ML364, the cells were collected and washed with PBS buffer and then stained with FITC-labelled anti-CD11b antibody (ab128797, Abcam, Shanghai, China) according to the manufacturer’s protocol. Differentiated cells (CD11b positive) were then determined by flow cytometer (BD Biosciences, San Diego, CA, USA). The results were analyzed using CytExpert 2.4 software.

### Immunofluorescence assay

Cells were fixed with 4% paraformaldehyde for 10 minutes at first. And then cells were permeabilized and blocked using QuickBlock™ blocking buffer for immunofluorescence staining (P0260, Beyotime, Shanghai, China) for 10 minutes. The cells were next incubated with the anti-PML antibody (dilution 1:50) and anti-USP2 antibody (dilution 1:100) overnight at 4 °C, followed by HRP-conjugated secondary antibody (dilution 1:200) for 2 hours at room temperature. The cell nuclei were stained with 4,6-diamidino-2-phenylindole (DAPI; C1002, Beyotime, Shanghai, China) before observing with the microscope. Confocal imaging was performed using a laser confocal microscope (Nikon, Nagoya, Japan).

### Proximity Ligation Assay (PLA)

Duolink® in situ Proximity Ligation Assay (DUO92101-1KT, Merck, Darmstadt, Germany) was used to detect interactions between USP2 and PML-RARα. The experiment was conducted following the manufacturer’s protocol.

### Statistical analysis

Data were obtained from three independent experiments and are expressed as mean ± standard deviation (SD). Statistical significance of differences between groups was determined by unpaired two-tailed Student’s t-test analysis and one-way analysis of variance (ANOVA). All graphs were analyzed using GraphPad Prism 8 software. The results were considered significant when *P* < 0.05 (n.s., *P* > 0.05; ∗*P* < 0.05; ∗∗*P* < 0.01; ∗∗∗*P* < 0.001 and ∗∗∗∗*P* < 0.0001).

## Results

### ML364 triggers PML/RARα degradation and shows a strong inhibitory effect on APL cells

To identify potential deubiquitinating enzymes that regulate the stability of PML-RARα, we treated NB4 cells with five reported DUB inhibitors (Supplementary Table 1) and examined the protein levels of PML-RARα. Among these compounds, ML364 significantly reduced the PML-RARα protein level (Figure 1A). To further validate this finding, we treated NB4 cells with varying doses of ML364 at different time points. The results demonstrated that ML364 decreased the protein level in a dose- and time-dependent manner (Figure 1B). Similar phenomena were observed in ATRA-resistant MR2 cells (Figure1C and Supplementary Figure S1).

**Figure 1.**
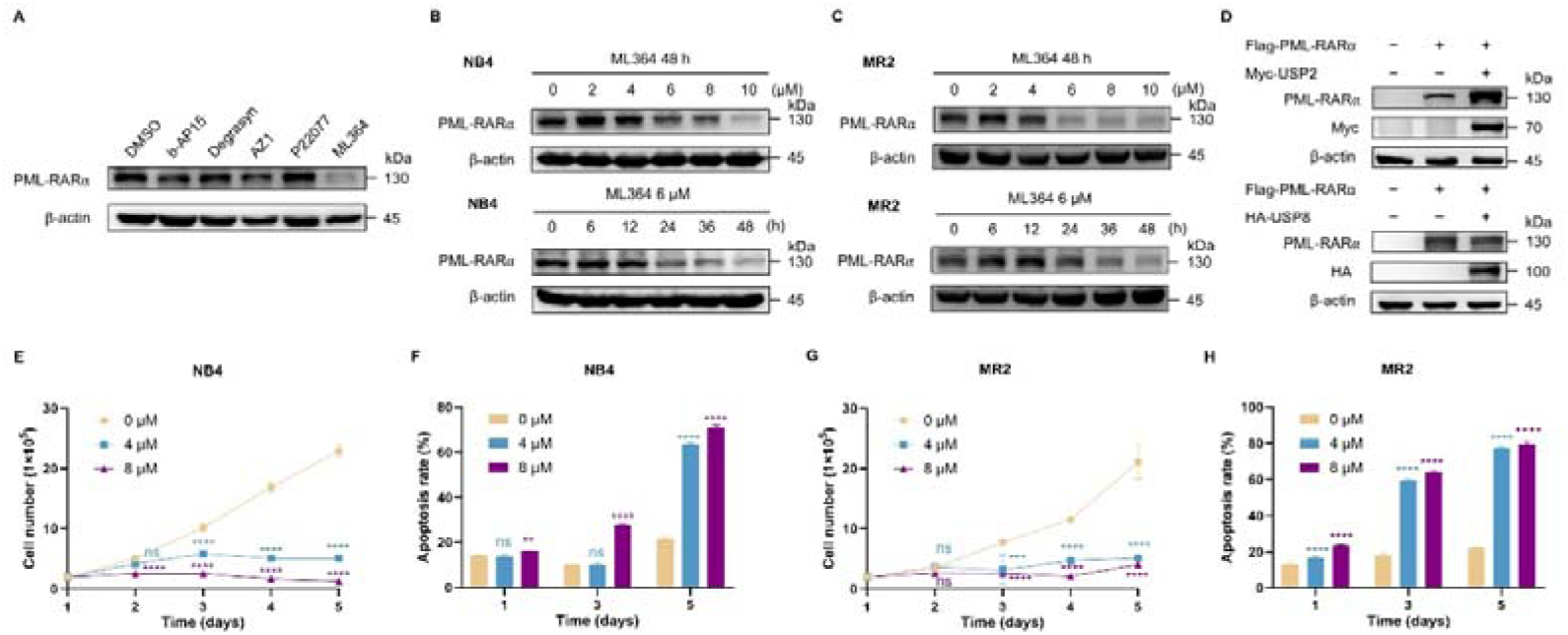
ML364 triggers PML/RARα degradation and shows a strong inhibitory effect on APL cells. (A) HEK293T cells stably expressing PML-RARα were treated with 5 DUB inhibitors at a respective IC_50_ concentration for 24 hours and the indicated proteins assessed by western blotting. All the western blot bands labeled as PML-RARα in this article were immunoblotted with anti-RARα antibody. (B) NB4 cells were exposed to varying concentrations of ML364 for 48 hours (top) or to 6 μM ML364 for the indicated times (bottom). (C) MR2 cells were exposed to varying concentrations of ML364 for 48 hours (top) or to 6 μM ML364 for the indicated times (bottom). (D) HEK293T cells were cotransfected with Flag-PML-RARα and Myc-USP2 or HA-USP8 plasmids, and the levels of exogenous PML-RARα were determined by western blotting. (E)(G) The proliferation of NB4 and MR2 cells exposed to the specified concentrations of ML364 was assessed using the trypan blue exclusion test for the indicated times. (F)(H) The apoptosis rate of NB4 and MR2 cells exposed to the specified concentrations of ML364 was assessed using flow cytometry analysis for the indicated times. Annexin V-positive cells were quantified with CytExpert software. (E-H) Data are presented as mean ± SD (n = 3); ∗*P* < 0.05, ∗∗*P* < 0.01, ∗∗∗*P* < 0.001 and ∗∗∗∗*P* < 0.0001 vs. 0 μM. The significance analysis was conducted by one-way ANOVA analysis.

ML364 has been previously reported as an inhibitor of USP2 and USP8 [19]. To determine which USP regulates PML-RARα expression, we overexpressed USP2 or USP8 in HEK293T cells transfected with exogenous PML-RARα (Figure 1D). Interestingly, the accumulation of PML-RARα protein was observed only in cells overexpressing USP2. These data suggest that targeting USP2 with ML364 can reduce PML-RARα protein levels.

Given the importance of PML-RARα for the survival of APL cells and the ability of ML364 to promote PML-RARα degradation by inhibiting USP2, we evaluated whether ML364 could inhibit APL cell proliferation and alleviate drug resistance. Treatment of NB4 cells with ML364 effectively suppressed proliferation and induced apoptosis (Figure 1E, F). Similarly, ML364 significantly reduced the proliferative capacity of MR2 cells (Figure 1G) and induced a substantial amount of apoptosis in these cells, too (Figure 1H). However, ML364 treatment did not alter the expression of CD11b, indicating that it did not induce differentiation in NB4 and MR2 cells (Supplementary Figure S2). These results suggest that ML364 may reduce the protein level of PML-RARα and inhibit the growth of APL cells by targeting USP2.

### USP2 knockdown decreases the protein levels of PML-RAR**α** and leads to effective APL eradication

To further elucidate the role of USP2 in acute promyelocytic leukemia (APL), we knocked down USP2 in NB4 cells and evaluated its effects on PML-RARα protein levels, cell proliferation, and cell death. Consistent with previous findings, USP2 knockdown led to a significant downregulation of PML-RARα protein levels (Figure 2A). Compared to the control group, USP2 knockdown markedly inhibited NB4 cell proliferation (Figure 2B) and induced significant apoptosis (Figure 2C and Supplementary Figure S3A). Similarly, USP2 knockdown in MR2 cells resulted in decreased PML-RARα protein levels (Figure 2D), inhibited proliferation (Figure 2E), and promoted apoptosis (Figure 2F and Supplementary Figure S3B). Conversely, overexpression of USP2 increased PML-RARα protein levels (Figure 2G) and enhanced NB4 cell proliferation (Figure 2H). These results collectively confirm that USP2 plays a crucial role in maintaining PML-RARα protein stability and promoting the proliferation of APL cells.

**Figure 2.**
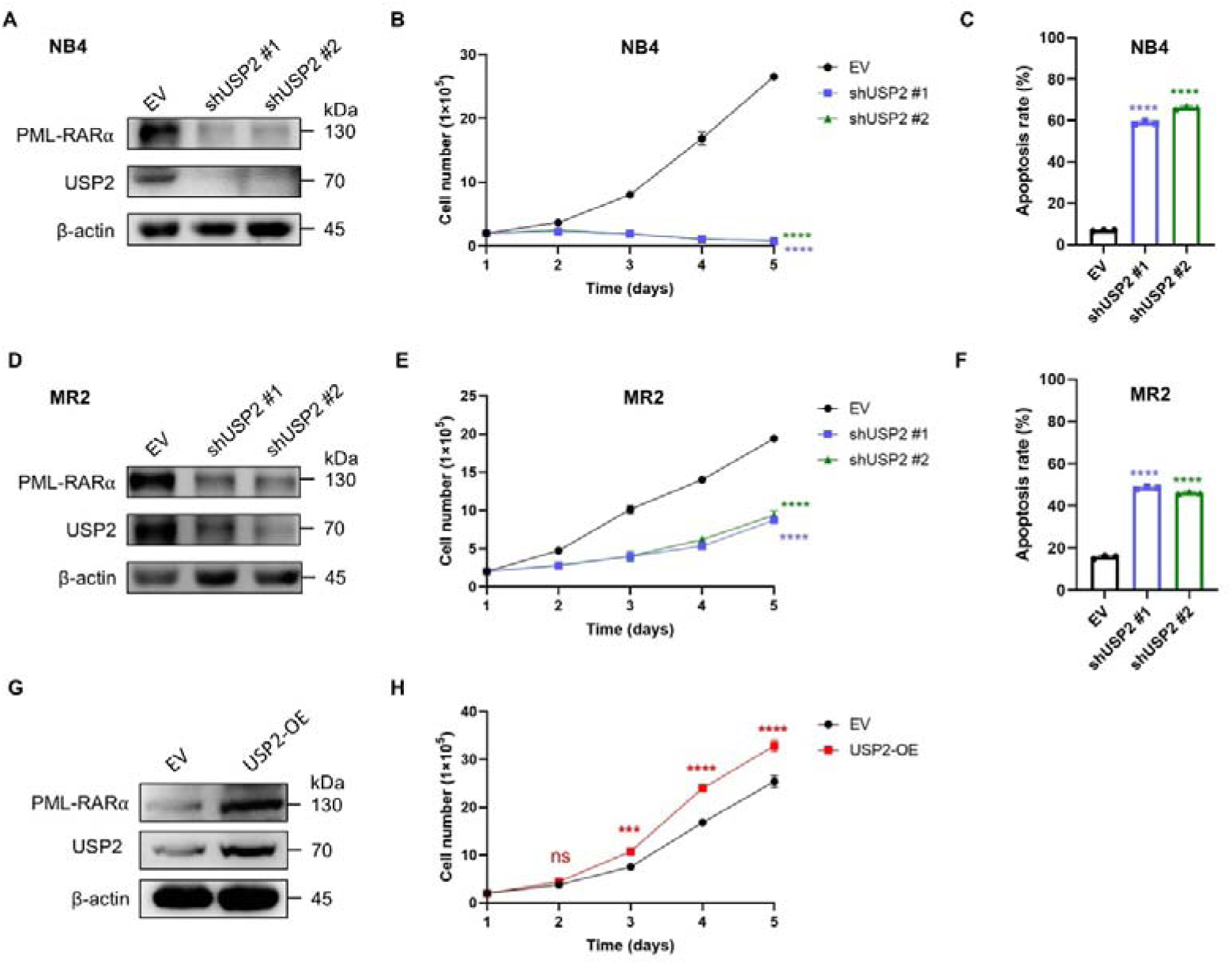
USP2 knockdown decreases the protein levels of PML-RARα and leads to effective APL eradication. (A-C) The expression of USP2 was silenced with shUSP2 #1 or shUSP2 #2 in NB4 cells. The indicated proteins were measured by western blotting (A). The proliferation of NB4 cells was assessed using the trypan blue exclusion test for the indicated times (B). The apoptosis rate was assessed using flow cytometry analysis (C). Annexin V-positive cells were quantified with CytExpert software. (D-F) USP2 was knocked down by shUSP2 #1 and shUSP2 #2 in MR2 cells. The indicated proteins were measured by western blotting (D). The proliferation of MR2 cells was assessed using the trypan blue exclusion test for the indicated times (E). The apoptosis rate was assessed using flow cytometry analysis for the indicated times (F). Annexin V-positive cells were quantified with CytExpert software. (G-H) USP2 was overexpressed in NB4 cells. The indicated proteins were measured by western blotting (G). The proliferation of NB4 cells with USP2 overexpression was assessed using the trypan blue exclusion test for the indicated times. (B)(C)(E)(F)(H) Data are presented as mean ± SD (n = 3); ∗*P* < 0.05, ∗∗*P* < 0.01, ∗∗∗*P* < 0.001 and ∗∗∗∗*P* < 0.0001 vs. EV. The significance analysis was conducted by one-way ANOVA analysis.

### USP2 regulates the stability of PML-RAR**α** through the ubiquitin-proteasome pathway

Given that ML364 inhibits the deubiquitinating activity of USP2, we hypothesized that ML364-induced downregulation of PML-RARα might be mediated through the ubiquitination-proteasome pathway. As expected, the proteasome inhibitor MG132, but not the autophagy inhibitor chloroquine (CQ), significantly prevented the decrease in PML-RARα levels induced by ML364 treatment (Figure 3A, B). Additionally, a cycloheximide (CHX) chase assay revealed that ML364 treatment accelerated PML-RARα degradation in NB4 cells (Figure 3C). These data indicate that ML364 reduces the protein level of PML-RARα by enhancing its proteasome-mediated degradation.

**Figure 3.**
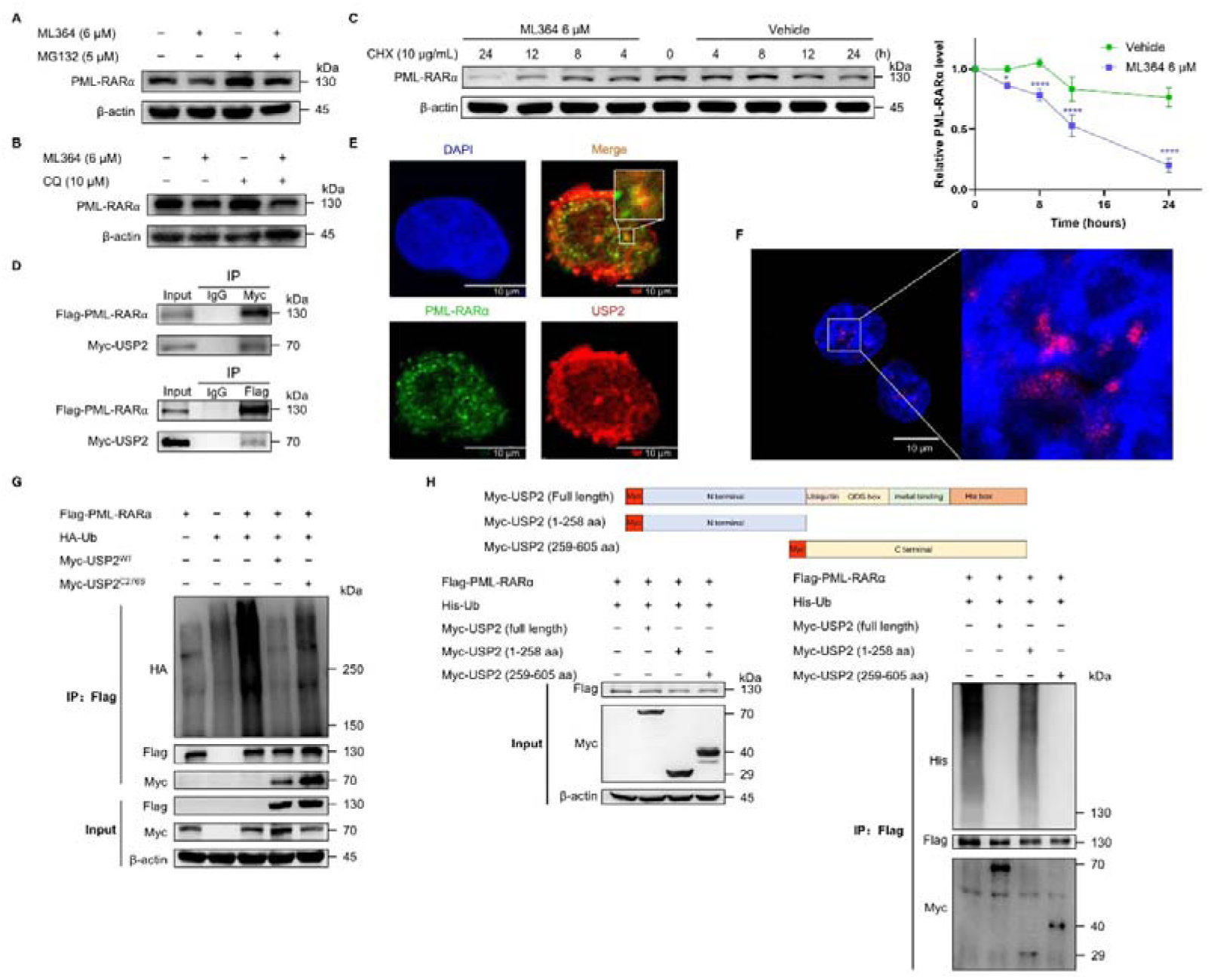
USP2 regulates the stability of PML-RARα through the ubiquitin-proteasome pathway. (A) NB4 cells were treated with 6 μM ML364 in the presence or absence of 5 μM MG132 for 24 hours. The protein levels were measured by western blotting. (B) NB4 cells were treated with 6 μM ML364 in the presence or absence of 10 μM CQ for 24 hours. The protein levels were measured by western blotting. (C) Left: NB4 cells were treated with CHX (10 μg/mL) in the presence or absence of ML364 (6 μM) for the indicated times. Right: Relative PML-RARα levels were analyzed using ImageJ. Data are presented as mean ± SD (n = 3); ∗*P* < 0.05, ∗∗*P* < 0.01, ∗∗∗*P* < 0.001 and ∗∗∗∗*P* < 0.0001 vs. Vehicle. The significance analysis was conducted by one-way ANOVA analysis. (D) HEK293T cells were cotransfected with Flag-PML-RARα and Myc-USP2. Cell lysates were immunoprecipitated and immunoblotted with antibodies against Flag and Myc. (E) Colocalization of PML-RARα and USP2 was detected by immunostaining in NB4 cells. Scale bar, 10 μm. (F) Interactions between PML-RARα and USP2 were detected by PLA in NB4 cells. Scale bar, 10 μm. (G) The deubiquitinating effect of USP2 on PML-RARα in cells. HEK293T cells were cotransfected with Flag-PML-RARα, His-Ub, Myc-USP2^WT^ and Myc-USP2^C276S^ as indicated. Cells were treated with MG132 (15 μM) for 6 hours before collected, and then immunoprecipitated with anti-Flag antibody. Ubiquitination of PML-RARα was detected by western blotting with anti-HA antibody. (H) Top: truncations of USP2 with Myc tag. Bottom: HEK293T cells were cotransfected with the indicated plasmids. Cells were treated with MG132 (15 μM) for 6 hours before collected, and then immunoprecipitated with anti-Flag antibody. Ubiquitination of PML-RARα was detected by western blotting with anti-His antibody.

To further confirm that targeting USP2 could regulate the ubiquitin-mediated degradation of PML-RARα, we first performed co-immunoprecipitation assays to examine the physical interaction between USP2 and PML-RARα. The results showed that Myc-USP2 interacts with Flag-PML-RARα (Figure 3D). Immunofluorescence assays and PLA experiments also demonstrated that USP2 and PML-RARα colocalize in APL cells (Figure 3E, F). Next, we investigated whether USP2 regulates the ubiquitination of PML-RARα. We transfected HEK293T cells with HA-Ub, Flag-PML-RARα, Myc-USP2WT, and its catalytic-site mutant, Myc-USP2C276S, and examined the ubiquitination of PML-RARα. The results showed that overexpression of USP2WT, but not USP2C276S, significantly reduced the ubiquitination level of PML-RARα (Figure 3G).

To map the specific domain of USP2 responsible for interacting with and regulating the stability of PML-RARα, we truncated USP2 into USP2 (1-258aa) and USP2 (259-605aa) and performed immunoprecipitation and western blot assays. The results indicated that the catalytic domain is crucial for USP2 to interact with and deubiquitinate PML-RARα (Figure 3H).

### USP2 also stabilizes different drug-resistant PML-RAR**α** mutants

Clinical observations have shown that the ligand-binding domain (LBD) of RARα and the PML-B2 domain are prone to mutations that confer resistance to all-trans retinoic acid (ATRA) and arsenic trioxide (ATO). We hypothesized that USP2 might also regulate the stability of these mutant proteins. To test this hypothesis, we transfected drug-resistant mutants, including ATO-resistant mutants (A216V and L218P) and ATRA-resistant mutants (R276Q and ΔF286) (Supplementary Figure S4), into HEK293T cells and treated them with varying concentrations of ML364. As shown in Figure 4A, ML364 effectively decreased the protein levels of these mutants.

**Figure 4.**
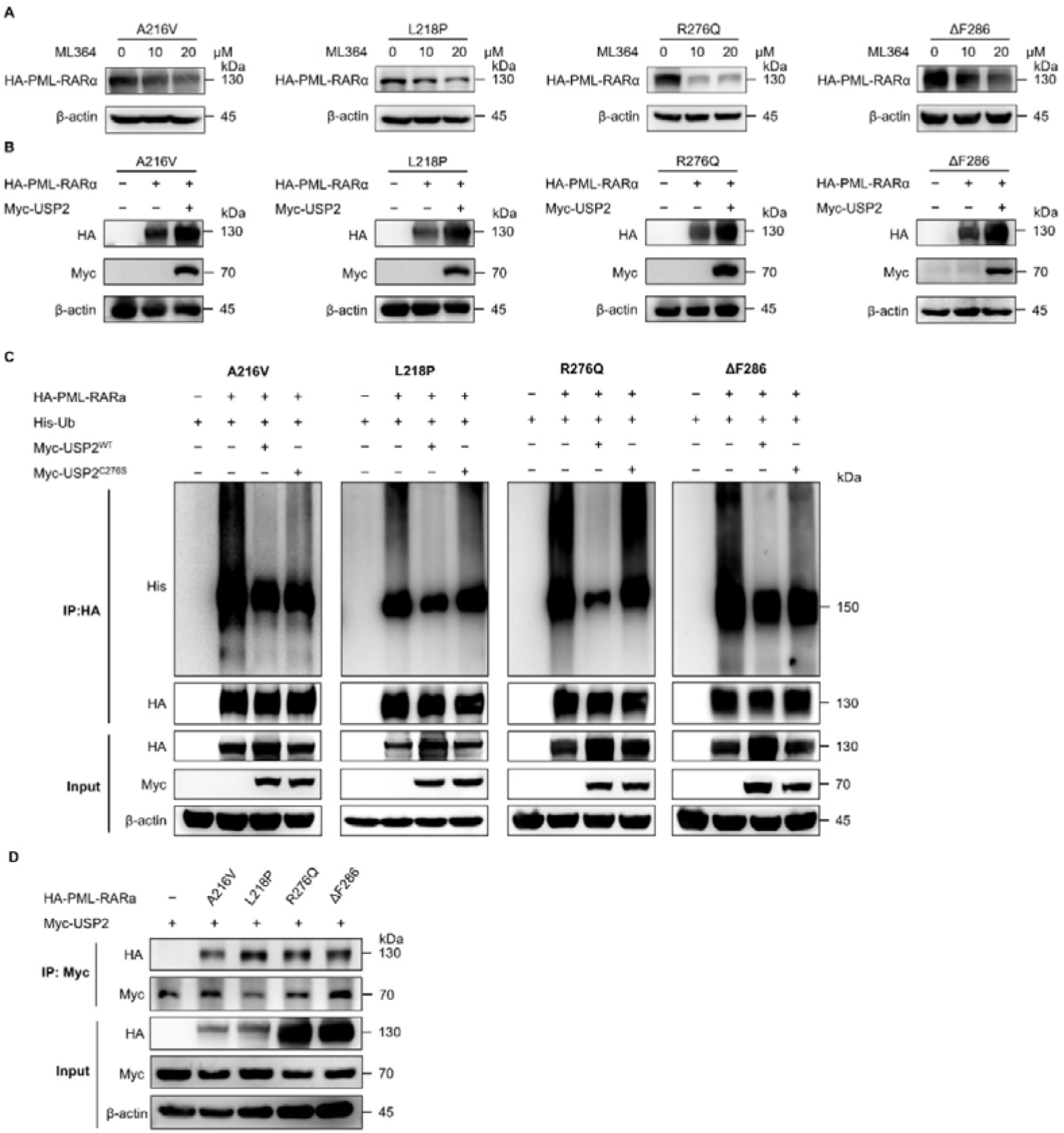
USP2 also stabilizes different drug-resistant PML-RARα mutants. (A) HEK293T cells were transfected with the indicated plasmids for 24 hours. Then cells were exposed to the indicated concentrations of ML364 for another 24 hours. Cell lysates were immunoblotted with anti-HA antibody. (B) HEK293T cells were cotransfected with Myc-USP2 and the indicated plasmids. Cell lysates were immunoblotted with anti-HA antibody. (C) HEK293T cells were transfected with the indicated plasmids. Cells were treated with MG132 (15 μM) for 6 hours before collected, and then immunoprecipitated with anti-HA antibody. Ubiquitination of PML-RARα was detected by western blotting with anti-His antibody. (D) The interactions between PML-RARα drug-resistant mutants (A216V, L218P, R276Q and ΔF286) and USP2 were detected by immunoprecipitation. HEK293T cells were transfected with the indicated plasmids. Cell lysates were immunoprecipitated with anti-Myc antibody and immunoblotted with anti-HA antibody.

To verify the role of USP2 in regulating the stability of PML-RARα mutants, we overexpressed USP2 alongside the aforementioned drug-resistant PML-RARα mutants. Interestingly, USP2 overexpression led to increased protein levels of each of the four resistant PML-RARα mutants (Figure 4B). Moreover, USP2 significantly reduced the ubiquitination levels of the resistant PML-RARα mutants, providing further evidence that USP2 may deubiquitinate PML-RARα (Figure 4C).

In reciprocal immunoprecipitation assays, overexpression of HA-tagged drug-resistant PML-RARα was detected in the immunocomplexes of Myc-USP2 (Figure 4D). These findings suggest that USP2 not only interacts with and regulates the stability of wild-type PML-RARα but also does so for mutant forms of PML-RARα.

### ML364 inhibits proliferation and promotes apoptosis of APL primary cells

To urther validate the effectiveness of ML364, we treated primary APL blasts with this compound. ML364 significantly inhibited cell proliferation in a dose-dependent manner (Figure 5A). After 72 hours of treatment, the IC_50_ values in five primary APL blasts ranged from 6.45 μM to 13.62 μM. ML364 also induced significant apoptosis in these cells, with the maximum apoptosis rate reaching 72.13% (Figure 5B). Furthermore, PML-RARα protein levels were significantly reduced in primary APL cells after ML364 treatment (Figure 5C). These results indicate that ML364 is effective in inhibiting the proliferation of primary APL cells.

**Figure 5.**
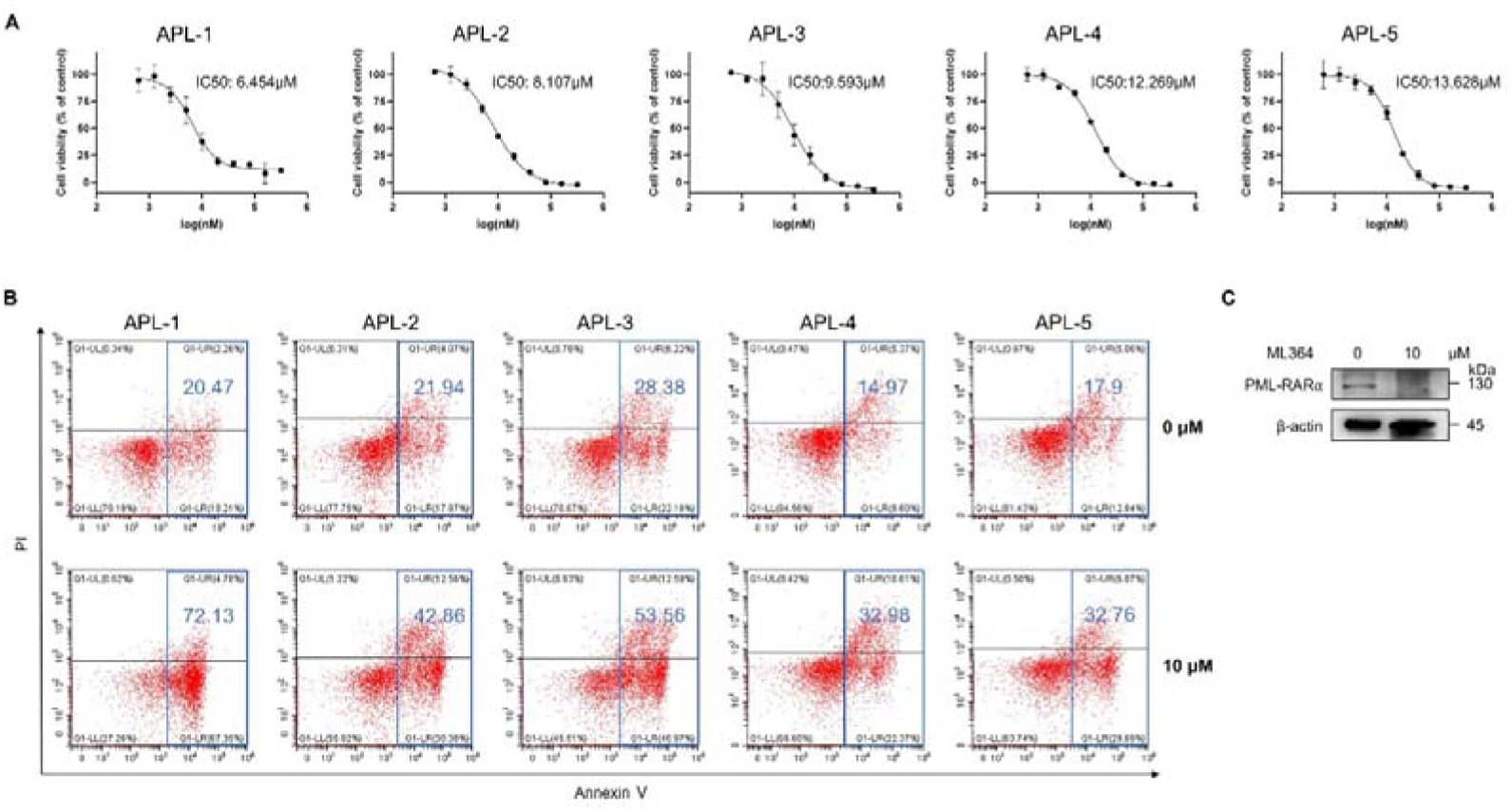
ML364 inhibits proliferation and promotes apoptosis of APL primary cells. (A) Primary APL cells derived from the bone marrow of 5 patients were treated with ML364 as the indicated concentrations for 72 hours. CCK-8 assay was used to analyze IC_50_ value of ML364 in primary APL cells. (B) The apoptosis rate of primary APL cells treated with or without ML364 (10 μM) for 72 hours. Annexin V-positive cells were quantified with CytExpert software. (C) Primary APL cells were treated with 10 μM ML364 for 72 hours.

## Discussion

Inducing the degradation of PML-RARα is a promising strategy to combat acute promyelocytic leukemia. In this study, we identified USP2 as a novel DUB for PML-RARα. USP2 stabilizes PML-RARα by removing its ubiquitin chains, thereby preventing its degradation. Inhibition of USP2 promotes PML-RARα degradation, including drug-resistant mutants. Our findings suggest that USP2 is a potential therapeutic target for overcoming PML-RARα mutation-induced resistance to ATRA and ATO.

The combination of ATRA and ATO synergistically induces PML-RARα degradation, leading to long-term remission in APL patients. However, a subset of patients develops resistance due to mutations in PML or RARα, which hinder ATRA or ATO binding, preventing their therapeutic effects. Therefore, identifying novel strategies to degrade PML-RARα is crucial. PML-RARα degradation is mediated through the ubiquitin-proteasome system, where E3 ubiquitin ligases add ubiquitin, and DUBs remove it. To promote PML-RARα degradation, we can either activate E3 ligases or inhibit DUBs. Since developing an E3 ligase activator is technically challenging, targeting DUBs with small-molecule inhibitors is a more practical approach. Identifying DUBs that regulate PML-RARα stability is therefore essential. Several DUBs, including YOD1 and USP37, have been reported to regulate PML-RARα or PLZF-RARα stability[18, 20]. In this study, we identified USP2 as a novel DUB for PML-RARα. Mechanistically, USP2 interacts with and removes ubiquitin from PML-RARα, leading to its stabilization. Conversely, USP2 inhibition promotes PML-RARα degradation. These findings confirm that USP2 is a bona fide DUB for PML-RARα.

A significant discovery in this study is that USP2 regulates not only wild-type PML-RARα but also its drug-resistant mutants. This suggests that targeting USP2 could overcome ATRA- and ATO-resistance in APL patients. Interestingly, USP2 also regulates the stability of PLZF-RARα, another pathogenic fusion protein associated with ATRA and ATO resistance in APL (Supplementary Figure S5A,B)[21, 22]. These findings highlight USP2 as a valuable therapeutic target for APL driven by RARα fusion proteins.

As a deubiquitinating enzyme, USP2 plays a broad role in cell proliferation, apoptosis, and metabolism [23–25]. It is known to be involved in multiple solid tumors, including prostate cancer, liver cancer, colorectal cancer, and breast cancer[26–28]. However, its role in hematological malignancies remains largely unexplored. Recently, we reported that USP2 is implicated in myeloma by regulating KRAS stability[29]. In this study, we provide further evidence that USP2 is a promising target for APL therapy. The USP2 inhibitor ML364 significantly inhibited cell proliferation and induced apoptosis in both ATRA-sensitive and ATRA-resistant APL cells, as well as in primary APL cells. ML364 represents a promising candidate for treating APL. USP2 has been reported to regulate multiple substrates, including Cyclin D1, MDM2, and KRAS, which could independently influence cell proliferation and apoptosis [19, 29, 30]. Our study shows that USP2 stabilizes PML-RARα, and its depletion reduces APL cell viability. However, we do not exclude the possibility that the degradation of other substrates, such as Cyclin D1, may also contribute to the observed anti-leukemic effects.In conclusion, we identified USP2 as a novel DUB that regulates the stability of PML-RARα and its drug-resistant mutants. Our findings suggest that targeting USP2 represents a potential therapeutic strategy for treating ATRA- and ATO-resistant APL.

