## Supplementary material for "Targeting USP2 induces degradation of PML-RARα with or without drug-resistant mutations in acute promyelocytic leukemia": Supplementary Information-0425-3.docx

**Supplementary Table 1.** The information and IC_50_ concentrations of 5 DUB inhibitors.

|  | **DUB inhibitor** | **CAS** | **Target** | **IC_50_ Concentration (μM)** |
| --- | --- | --- | --- | --- |
| 1 | b-AP15 | 1009817-63-3 | USP14, UCHL5 | 1 |
| 2 | Degrasyn (WP1130) | 856243-80-6 | USP9x, USP5, USP14, UCH37 | 1 |
| 3 | USP25/28 inhibitor AZ1 | 2165322-94-9 | USP25/28 | 2 |
| 4 | P22077 | 1247819-59-5 | USP7, USP47 | 10 |
| 5 | ML364 | 1991986-30-1 | USP2, USP8 | 10 |

**Supplementary figures.**

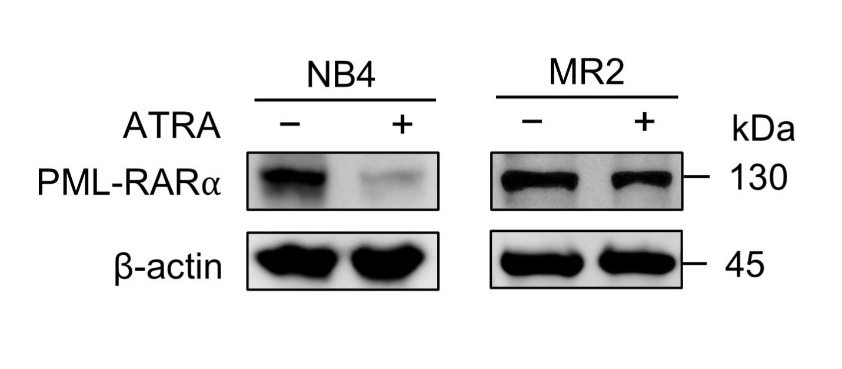

**Supplementary Figure S1. Verification of ATRA resistance in MR2 cells.** NB4 and MR2 cells were treated with 1 μM ATRA for 24 hours and the indicated proteins were assessed by western blotting.

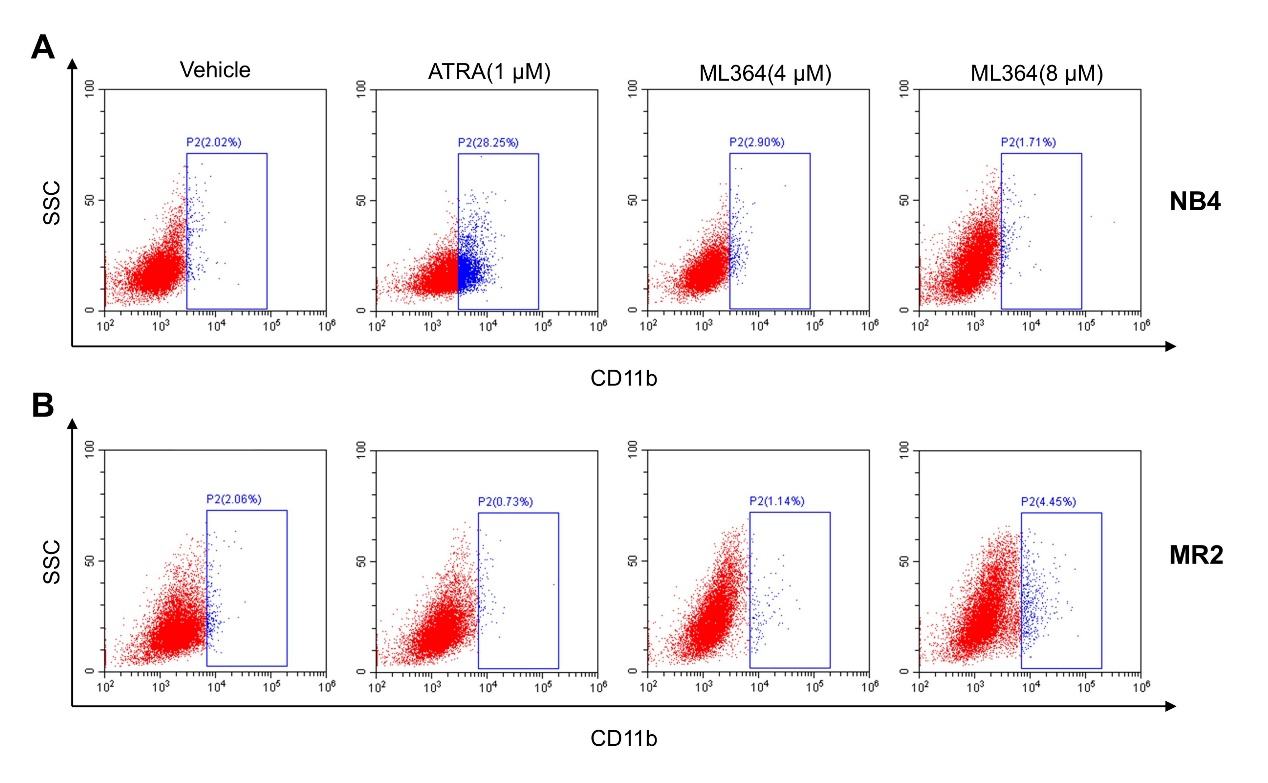

**Supplementary Figure S2. The effect of ML364 on cell differentiation.** NB4 (A) and MR2 (B) cells were treated with ATRA and ML364 as the indicated concentrations for 72 hours. Flow cytometry was used to analyze whether cells has been induced differentiation. CD11b-positive cells were quantified with CytExpert software.

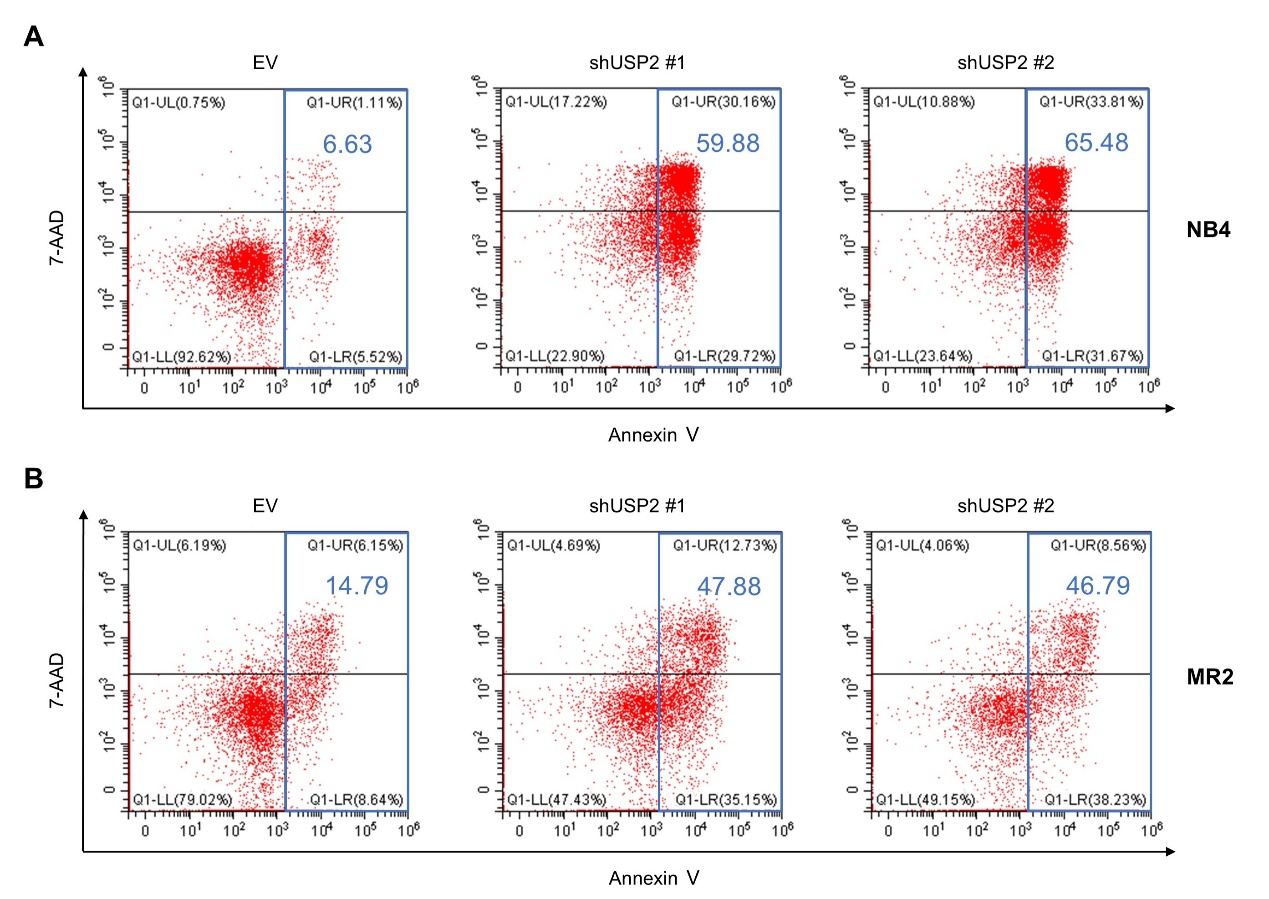

**Supplementary Figure S3. Knockdown of USP2 leads to apoptosis in APL cells.** Knockdown of USP2 was performed using shUSP2 #1 or shUSP2 #2. The apoptosis rate of NB4 (A) and MR2 (B) cells was assessed using flow cytometry analysis. Annexin V-positive cells were quantified with CytExpert software.

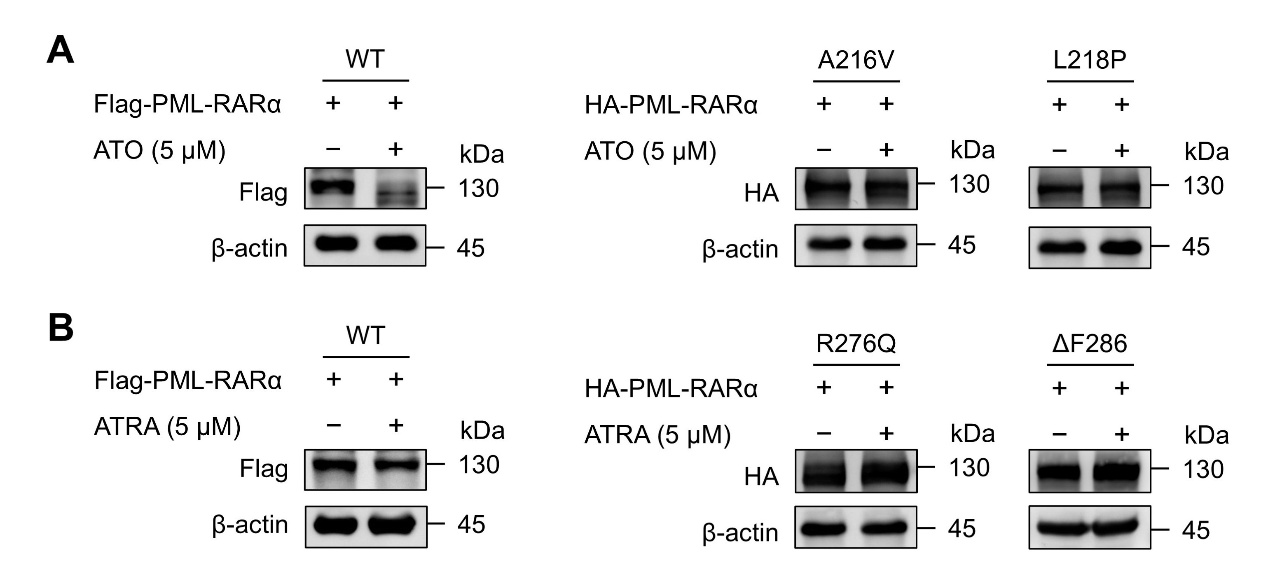

**Supplementary Figure S4. Validation of PML-RARα drug-resistant mutants (A216V, L218P, R276Q and ΔF286).** HEK293T cells were transfected with the indicated plasmids for 24 hours. Then cells were exposed to ATO (A) or ATRA (B) for another 24 hours. Cell lysates were immunoblotted with anti-Flag or anti-HA antibody.

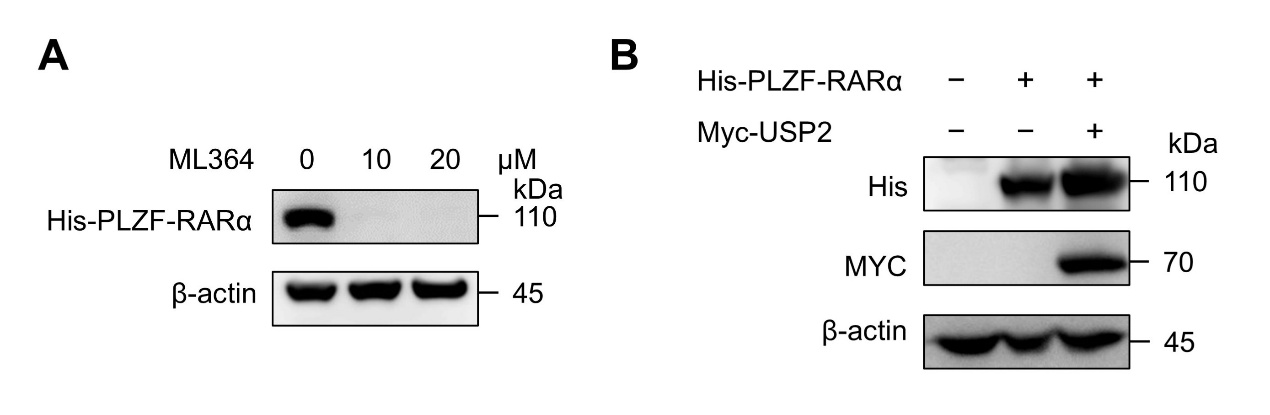

**Supplementary Figure S5. Effect of ML364 and USP2 overexpression on PLZF-RARα.** (A) HEK293T cells were transfected with His-PLZF-RARα for 24 hours. Then cells were exposed to the indicated concentrations of ML364 for another 24 hours. Cell lysates were immunoblotted with anti-His antibody. (B) Effect of USP2 overexpression on the protein level of PLZF- RARα. HEK293T cells were transfected with Myc-USP2 and His-PLZF-RARα.
